# Visualization of a Limonene Synthesis Metabolon inside Living Bacteria by Hyperspectral SRS Microscopy

**DOI:** 10.1101/2022.06.30.498009

**Authors:** Jing Zhang, Jonghyeon Shin, Nathan Tague, Haonan Lin, Meng Zhang, Xiaowei Ge, Wilson Wong, Mary J. Dunlop, Ji-Xin Cheng

## Abstract

Metabolons consisting of cellular structure elements and sequential metabolic enzymes are expected to be involved in diverse biological processes. However, direct visualization of metabolons in prokaryotic cells is still challenging. In this study, we report direct visualization of concentrated subcellular regions of limonene synthesis inside single engineered *Escherichia coli* by using hyperspectral stimulated Raman scattering (hSRS) microscopy. Equipped with spectral unmixing, hSRS imaging provides a reliable method to quantify intracellular limonene content. In *E. coli* strains with a complete limonene synthesis pathway, intracellular limonene is found locally concentrated and colocalized with proteins. Furthermore, dual-modality SRS and two-photon fluorescence imaging showed colocalization of limonene and GFP-fused limonene synthase.

**Significance Statement:** Monitoring biosynthesis activity at the single-cell level is key to metabolic engineering but is still difficult to achieve in a label-free manner. Using hyperspectral stimulated Raman scattering imaging in the 670-900 cm^−1^ region, we visualized localized limonene synthesis inside engineered *E. coli*. The colocalization of limonene and GFP-fused limonene synthase was confirmed by co-registered stimulated Raman scattering and two-photon fluorescence images. Our finding suggests a limonene synthesis metabolon with a polar distribution inside the cells. This finding expands our knowledge of *de novo* limonene biosynthesis in engineered bacteria and highlights the potential of SRS chemical imaging in metabolic engineering research.

## Introduction

A metabolon is a supramolecular complex of sequential metabolic enzymes (1, 2). Metabolons provide multiple advantages for enhanced product flux, and protection from toxic metabolic intermediates (3, 4). Moreover, a few examples of metabolons such as a glucosome and purinosome have been reported (5–7). Advanced imaging techniques such as gas cluster ion beam secondary ion mass Spectrometry (GCIB-SIMS), super-resolution fluorescence microscopy, and fluorescence resonance energy transfer (FRET) microscopy have been employed to image metabolons, e.g. purinosomes in eukaryotic cells (8–10). However, the investigation of metabolons in a single bacterial cell remains extremely challenging, especially when the metabolites involved are small molecules for which fluorescent reporters are not available.

Limonene is a type of cyclic monoterpenes that widely exist in plants. Both limonene and its derivatives have diverse applications in the food industry and pharmaceuticals (11–13). Despite increasing demand, the supply of limonene, mainly produced from food processing, is unstable (14). Empowered by the latest advancements in genetic engineering and metabolic engineering, microbial fermentation facilitating the conversion of limonene from low-value materials like glucose has been a promising approach for the bulk and stable production of limonene (15, 16). However, cell-to-cell variation of biochemical synthesis hinders the maximum yield and the lack of detailed single-cell level characterization is one of the bottlenecks to solve the problem.

Recently, spontaneous Raman has been applied to study metabolite production in engineered fungi or bacteria (17, 18). Due to the small Raman cross-sections of biological samples, the speed of spontaneous Raman is limited, and it usually requires tens of milliseconds to seconds dwell time. This long acquisition time hinders the further utility of spontaneous Raman in biological applications.

Employing a two-laser system (a pump beam and a Stokes beam), stimulated Raman scattering (SRS) offers accelerated imaging speed compared to spontaneous Raman (19–21). The rapid identification of Raman signals offered by SRS paves the way for single-cell imaging with live-cell compatibility (22–24). Compared to fluorescence imaging, SRS has the potential to detect molecules that do not have commercially available fluorescent probes. Here, we report direct visualization of localized limonene synthesis inside single engineered *Escherichia coli* by hyperspectral stimulated Raman scattering (hSRS). Compared to previous study of limonene biosynthesis detection by single-color SRS (25, 26), hSRS allows simultaneous quantification of multiple chemicals inside one specimen by recording a Raman spectrum at each pixel. Intracellular compositions of various biological specimens have been studied using hSRS, ranging from tissues (27, 28), to cancerous cells (29, 30), to bacteria (31, 32). Many of these applications employ the C-H stretching region (2800 – 3100 cm^−1^) for its strong SRS signal or the C-D stretching region (2070 – 2300 cm^−1^) for selective imaging of Raman tags. However, both C-H region and C-D band in the silent region lack specificity in detecting small-molecule synthesis because of Raman signal contributions from other intracellular biomacromolecules. In comparison, the fingerprint region (600-1800 cm^−1^) takes advantage of the rich spectral features of limonene (33), making hSRS in the fingerprint region an ideal tool to study limonene biosynthesis at the single-cell level.

In this study, we harness polygon-scanner-based hSRS to map the *de novo* synthesized limonene distribution inside *E. coli* cells. We first verify that limonene is synthesized from engineered *E. coli* using gas chromatography-mass spectrometry (GC-MS). To investigate subcellular limonene distribution, L_1_-regularized least squares fitting was used for hSRS image stack in 670 – 900 cm^−1^ region, enabling selective mapping of limonene and proteins inside individual *E. coli* cells. The chemical content and spatial distribution of the limonene-rich aggregates were then characterized by single-cell and single-aggregate segmentation. In addition, two-photon fluorescence (TPF) imaging of GFP-fused limonene synthase supports the colocalization of limonene with limonene synthase. Together, our data support the possible existence of metabolons in engineered *E. coli*. The result also highlights the potential of hSRS imaging in metabolic engineering through multiplexed biomolecular analysis at the sub-cellular level.

## Results

### Limonene-producing *E. coli* strains and mass spectrometry measurements

We harnessed the heterogeneous mevalonate pathway to produce limonene in *E. coli* (**Figure 1a**) (34). Isoprenoid precursors, isopentenyl pyrophosphate (IPP) and dimenthylallyl pyrophosphate (DMAPP), are products of the mevalonate pathway derived from acetyl-CoA, one of the key node molecules in *E. coli* central metabolism. A geranyl pyrophosphate synthase (GPPS) converts IPP and DMAPP to geranyl pyrophosphate (GPP) which is a universal precursor of terpenes and a limonene synthase (LS) specifically converts GPP to limonene. For easy genetic modification, we distributed necessary enzymes in two plasmids: one plasmid encodes the mevalonate pathway and the other encodes GPPS and LS which are translationally fused (denoted as a two-plasmid system, **Figure 1b**) (34). *E. coli* strain BW25113 was used as the host. Limonene production is induced by a small molecule isopropyl β-d-1-thiogalactopyranoside (IPTG). IPTG is a molecular reagent that induces expression where the gene is under the control of the lac repressor. We then used gas chromatography–mass spectrometry (GC-MS) to confirm the successful limonene synthesis in these engineered strains (**Figure 1c**).

**Figure 1.**
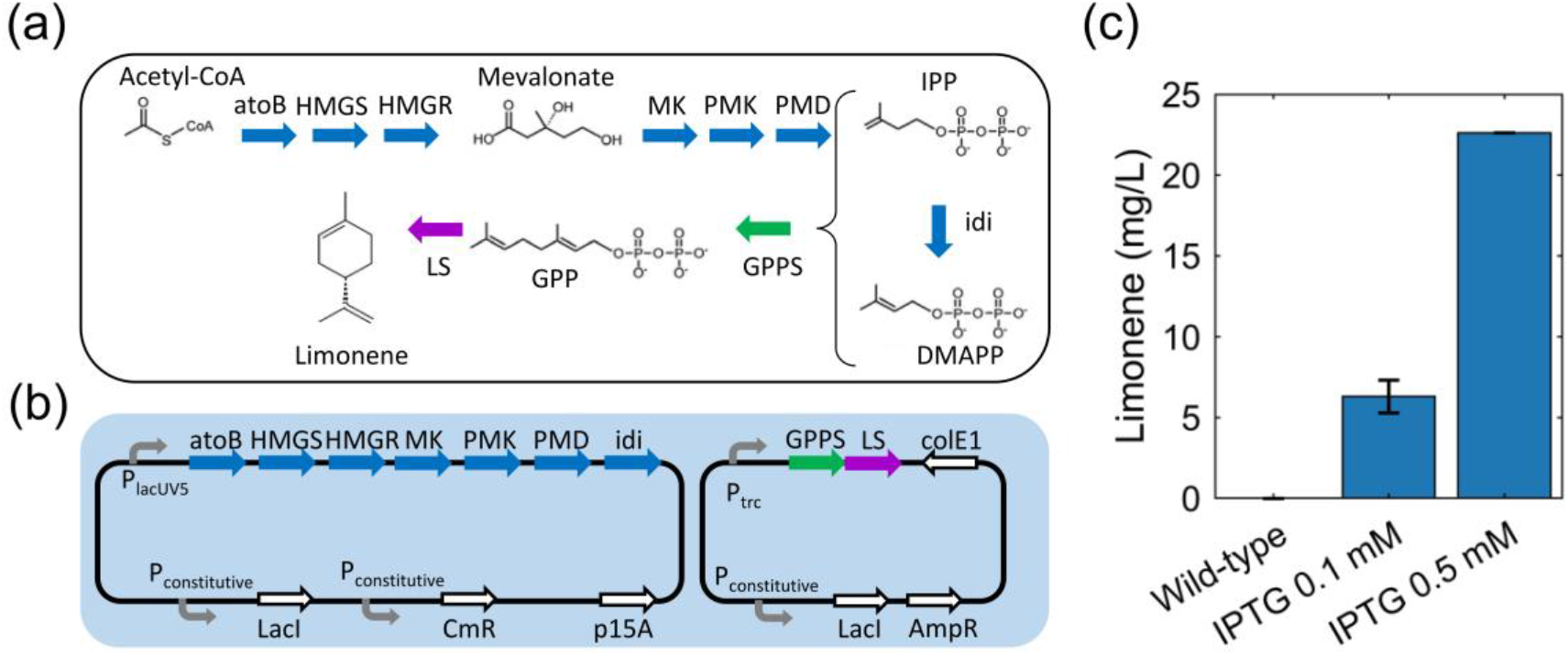
Limonene-producing *E. coli* strains and GC-MS measurements. (a) Metabolic pathway from Acetyl-CoA to limonene in *E. coli*. Enzymes and some of the reaction intermediates necessary for the production of limonene are shown. AtoB: acetoacetyl-CoA synthase; HMGS: HMG-CoA synthase; HMGR: MG-CoA reductase; MK: Mevalonate Kinase; PMK: PhosphoMevalonate Kinase; PMD: PhosphoMevalonate Decarboxylase; IPP: isopentenyl pyrophosphate; idi: isopentenyl diphosphate isomerase; DMAPP: dimenthylallyl pyrophosphate; GPPS: Geranyl PyroPhosphate Synthase; GPP: geranyl pyrophosphate; LS: limonene synthase. (b) Genetic design of limonene pathway for strains with a two-plasmid system. (c) GC-MS measurements of microbial production of limonene (BW25113 strain with a two-plasmid system). Error bar: std.

### Hyperspectral SRS imaging unveils chemical aggregates inside limonene-producing *E. coli*

To visualize limonene biosynthesis at the single-cell level, we utilized both Raman spectroscopy and SRS microscopy. Raman spectroscopy was used to determine the vibrational signatures of limonene and its precursors (**Figure 2a,b**). Pure limonene, glycerol trioleate (GT), and bovine serum albumin (BSA) were measured using a LabRAM HR800 confocal Raman microscopy. Spectra of geraniol, prenol, and isoprenol from (35) were used to present the spectra of geranyl diphosphate (GPP), dimethylallyl diphosphate (DMAPP), and isopentenyl diphosphate (IPP), which are the intermediates in the limonene synthesis pathway. Limonene exhibits strong Raman signals in the high wavenumber C-H stretching region (2800 to 3100 cm^−1^), but this region lacks limonene specificity as C-H bonds are the most abundant functional groups in the cell. In the fingerprint region (600 to 1800 cm^−1^), limonene shows several signature peaks. We chose the peak at 760 cm^−1^ considering the high intensity of limonene and the relatively low background contributed by other biomolecules in the 670 to 900 cm^−1^ region. This unique limonene peak stems from a combination of ring and CH2 vibration in limonene (36). We also measured spectra of GT and BSA to confirm that neither protein nor fatty acids inside the cells do not have a significant contribution to the 760 cm^−1^ Raman peak.

**Figure 2.**
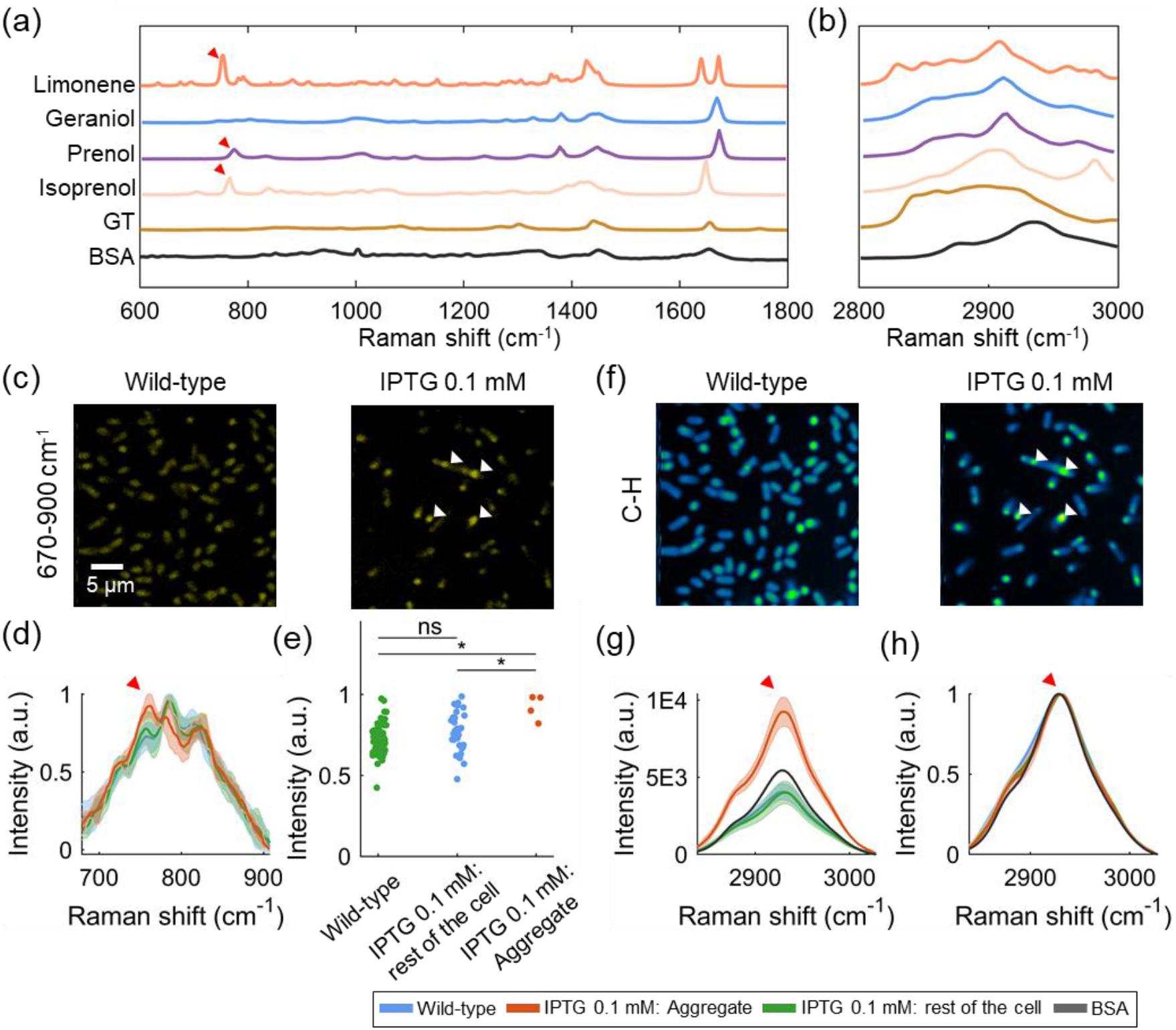
Hyperspectral SRS imaging unveils chemical aggregates inside limonene-producing *E. coli*. (a) Spontaneous Raman spectra of limonene, geraniol, prenol, isoprenol, GT (glycerol trioleate), and BSA (bovine serum albumin) in the fingerprint window (600-1800 cm^−1^). Spectra of geraniol, prenol, and isoprenol represent the spectra of precursors in limonene synthesis pathway (GPP, DMAPP, IPP). (b) Spontaneous Raman spectra of limonene, geraniol, prenol, isoprenol, GT, and BSA in the C-H window. The spectra of GT and BSA were included to show that neither protein nor fatty acids inside the cells have a significant contribution to the 760 cm^−1^ Raman peak. Red arrows denote peaks at Raman shift 760 cm^−1^ (limonene), 774 cm^−1^ (prenol), and 767 cm^−1^ (isoprenol). (c) Spectrally summed hSRS images of BW25113 strain (fingerprint window). Left: wild-type; Right: limonene-producing strain. White arrows denote the position of the intracellular aggregates. (d) Average of the normalized spectra of the wild-type strain, aggregate of limonene-producing strain, and non-aggregating part of limonene-producing strain. Red arrow marks 760 cm^−1^ Raman peak. Shaded error bar: std. (e) Scatter plot of the Raman intensity at 760 cm^−1^ per region of the normalized spectra in (d). (f) Spectrally summed hSRS images of BW25113 strain (C-H window). Left: wild-type; Right: limonene-producing strain. (g) Average of the raw spectra of the wild-type strain, aggregates of limonene-producing strain, and non-aggregating part of limonene-producing strain. Red arrow marks 2930 cm^−1^ Raman peak. (h) Average of the normalized spectra of wild-type strain, aggregates of limonene-producing strain, and non-aggregating part of limonene-producing strain. Red arrow marks 2930 cm^−1^ Raman peak. Shaded error bar: std.

Next, hSRS imaging was utilized to characterize the spatial distribution of limonene inside single engineered *E. coli* cells. We employed an ultrafast delay-tuning hSRS scheme (22) shown in **Figure S1**. Briefly, polygon-scanner-based SRS collects image stacks in a “Λ-x-y” manner and provides an acquisition speed of down to 20 μs per spectrum with a 10 cm^−1^ resolution in the fingerprint region (22). In this study, each single spectrum in the SRS stack takes 120 μs, which improved the spectral fidelity and eliminated low-frequency noise compared to the conventional motorized-stage-based hSRS scheme. It thus provides great advantages for differentiating limonene from other biomolecules inside cells. **Figure S2** shows the linearity and sensitivity of limonene detection at the 760 cm^−1^ Raman peak. The detection limit is found to be ~21 mM at a speed of 0.89 seconds per frame of 200×200 pixels. Compared to the conventional frame-by-frame scheme, the use of the polygon scanner increased the hSRS imaging speed by about 5 times at the same signal-to-noise ratio (SNR) (**Figure S3**).

By hSRS imaging, intracellular aggregates were observed in the 670-900 cm^−1^ region for the limonene-producing strains, as indicated by arrows in **Figure 2c**. In contrast, the control strains without the heterologous limonene synthesis pathway did not exhibit obvious intracellular aggregates. **Figure 2d** shows the fingerprint Raman spectra of the aggregates and the rest of the cell. Specifically, the Raman peak at 760 cm^−1^ confirms the presence of limonene inside these aggregates. The spectrum of the rest of the cell, lacking the 760 cm^−1^ Raman peak, is comparable to the spectrum of the wild-type strain (Wilcoxon rank sum test, **Figure 2e**). As a comparison, hSRS imaging in the C-H region was also performed. Intracellular aggregates were also observed in the limonene-producing strain but not in the control strain (**Figure 2f**). As opposed to the fingerprint region, the spectra in the C-H region do not distinguish the aggregate from the rest of the cell, as the C-H region does not have enough specificity for limonene detection (**Figure 2g,h**). Besides, in the C-H region, the spectra of aggregates resemble the spectrum of BSA but not limonene, indicating the coexistence of proteins with the locally aggregated limonene. Because of the low solubility of limonene and the intermediates in water, these proteins might offer protection to hydrophobic molecules against the hydrophilic environment of the cytoplasm. Further testing of the chemical composition and potential functions of these intracellular aggregates is shown in the following sections.

### Pixel-wise spectral unmixing reveals the chemical composition of the intracellular aggregates

To extract the limonene distribution map from the hyperspectral stack, we applied the least absolute shrinkage and selection operator (LASSO) (37). The effectiveness of LASSO in unmixing hSRS images in the fingerprint region has been tested on various biological samples (22). Compared to least squares fitting, LASSO introduces a pixel-wise sparsity constraint. It hypothesizes that each pixel of our hyperspectral image is dominated by sparsity components. This feature makes LASSO ideal for our study because limonene is localized in the intracellular aggregates and the rest of the cell (the non-aggregating part) has the same spectral profile as the control strain (**Figure 2d**). Therefore, using the reference spectra for pure limonene, wild-type strain, and background (of each hSRS image stack), chemical maps of three channels can be generated, denoted as limonene, cell body, and background.

By LASSO analysis of the fingerprint hSRS data (**Figure 3a**), we generated concentration maps of limonene (**Figure 3b**). Locally concentrated limonene was found in engineered strains but not in the wild-type. The cell body channel was similar across all samples (**Figure 3c**). By single-cell and single-aggregate segmentation (**Figure S4**, details in the Methods section), segmentation maps of two types of cell regions were generated: aggregates and the rest of the cell. By averaging the limonene intensity of both regions, the limonene amount of each aggregate and the rest of the cell is plotted (**Figure 3d**). This result indicates the limonene intensity of aggregate depends on IPTG concentration, which is qualitatively correlated to GC-MS measurements (**Figure 1c**). Importantly, SRS imaging and LASSO analysis quantitatively show that the produced limonene is locally concentrated and not spread over the whole cell.

**Figure 3.**
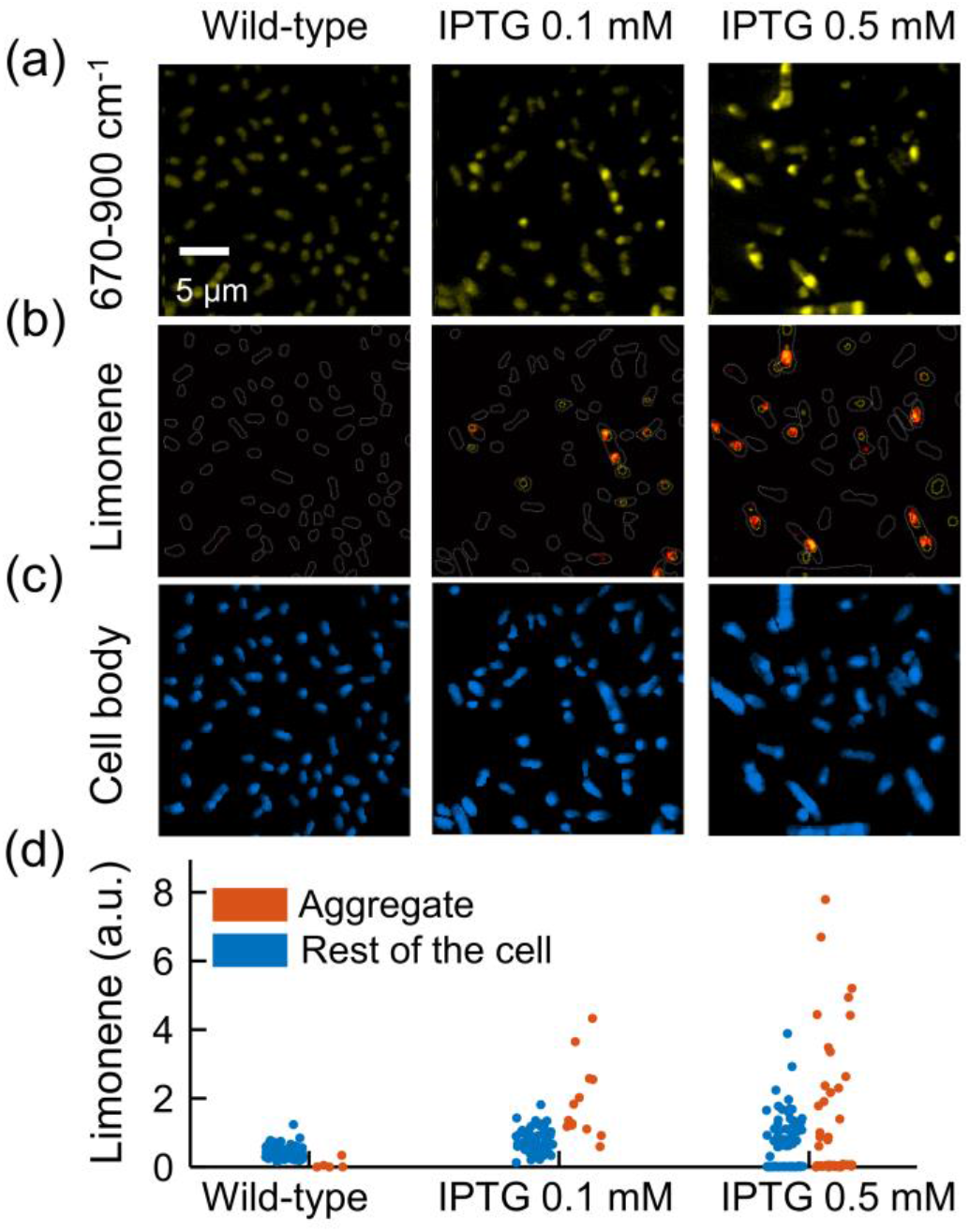
Chemical composition analysis of limonene-producing *E. coli* strains. (a) Spectrally summed hSRS images (670-900 cm^−1^). Left: wild-type; middle: Limonene-producing strain with 0.1 mM IPTG; right: Limonene-producing strain with 0.5 mM IPTG. (b) Chemical map of limonene. Localized limonene distribution is observed in limonene-producing strain. White lines: contour of each single cell. Yellow lines: contour of each single aggregate. (c) Chemical map of cell body. (d) Scatter plot of limonene intensity in aggregates and rest of the cell. Each dot represents the averaged limonene intensity in a single aggregate.

To verify that the SRS-mapped limonene indeed originates from *de novo* synthesis, we tested *E. coli* strains with incomplete synthesis pathways in addition to the wild-type strain and the strain with the complete limonene synthesis pathway. Specifically, we tested three incomplete pathways: ΔGPPS without GPPS but having all other enzymes, ΔLS without LS but having all other enzymes, and GPPS-LS only having GPPS and LS but none of the upstream genes in the mevalonate pathway. Based on hSRS images in the 670-900 cm^−1^ spectral region (**Figure 4a**), the concentration map of limonene was produced via LASSO analysis (**Figure 4b**). A high abundance of limonene was only observed in the strain with a full limonene synthesis pathway, in the form of aggregates. By applying single-aggregate segmentation to SRS images (C-H region, **Figure 4c**) of the same specimens, limonene-channel intensities in aggregate and the rest of the cell are plotted in **Figure 4d**. Using a threshold sweeping method (**Figure S5**), 1.1 was set as the limonene intensity threshold (red dashed line in **Figure 4d**) to distinguish the limonene-rich aggregates. Using this threshold, the percentage of cells with limonene-rich aggregates is 16.9% in the strain with a complete pathway. Further tests with a different construct design are shown in the following section.

**Figure 4.**
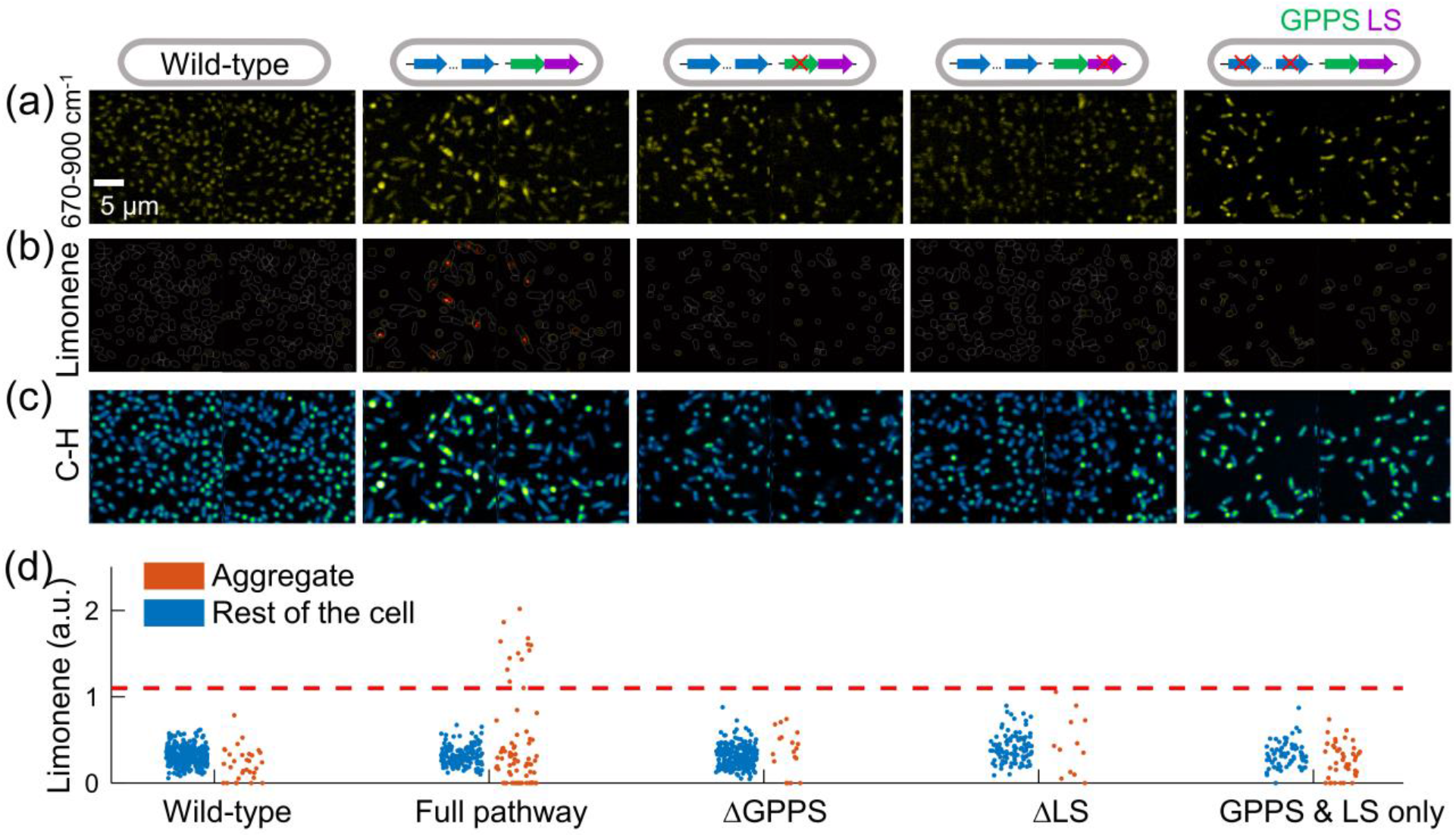
Characterization of limonene distribution in *E. coli* with complete or incomplete limonene synthesis pathways. The genetic design of each strain is shown in the top row. (a) Spectrally summed hSRS images in the fingerprint region. Aggregates are observed in limonene-producing *E. coli* strains. (b) Chemical maps of limonene. Localized limonene distribution is observed in limonene-producing strain. White lines: contour of each single cell. Yellow lines: contour of each single aggregate. (c) Spectrally summed SRS images at C-H region. (d) Scatter plot of limonene intensity in aggregates and rest of the cell. Red dashed line denoted the threshold of limonene-rich aggregates.

### Localized limonene synthesis is confirmed in in one-plasmid limonene-producing strains

To confirm the evidence of localized limonene aggregate formation, engineered strains with a one-plasmid version of the limonene production pathway were tested as a different construct design (denoted as one-plasmid system, **Figure 5a**). As production is impacted by the strain background, we tested *E. coli* strain DH1 in addition to BW25113. These one-plasmid strains show higher limonene production in GC-MS measurements than the two-plasmid design in BW25113 (**Figure S6a,b**). To further demonstrate the *in situ* live-cell imaging compatibility of our imaging system, we directly grew the cells on a thin layer of agarose gel pads after IPTG induction (method). With growth medium and air exchange provided by the agarose pad (38), colonies composed of a single layer of cells form on top of the agarose pad. After culturing for 24 hours, the cells growing on the agarose pad were directly imaged using SRS. Using this protocol, we tested the *E. coli* DH1 strain with one-plasmid, and we were also able to observe intracellular aggregates in both the C-H and the 670-900 cm^−1^ region (**Figure 5b,c**). Limonene concentration maps were produced using LASSO analysis (**Figure 5d**), showing that limonene is localized and enriched in these aggregates. By segmenting these bright spots from the rest of the cell, fingerprint spectra of the aggregates were extracted and exhibited a distinct Raman peak at 760 cm^−1^ unique to limonene (**Figure 5e**). Similar results were obtained with another host strain (**Figure S6c-f**).

**Figure 5.**
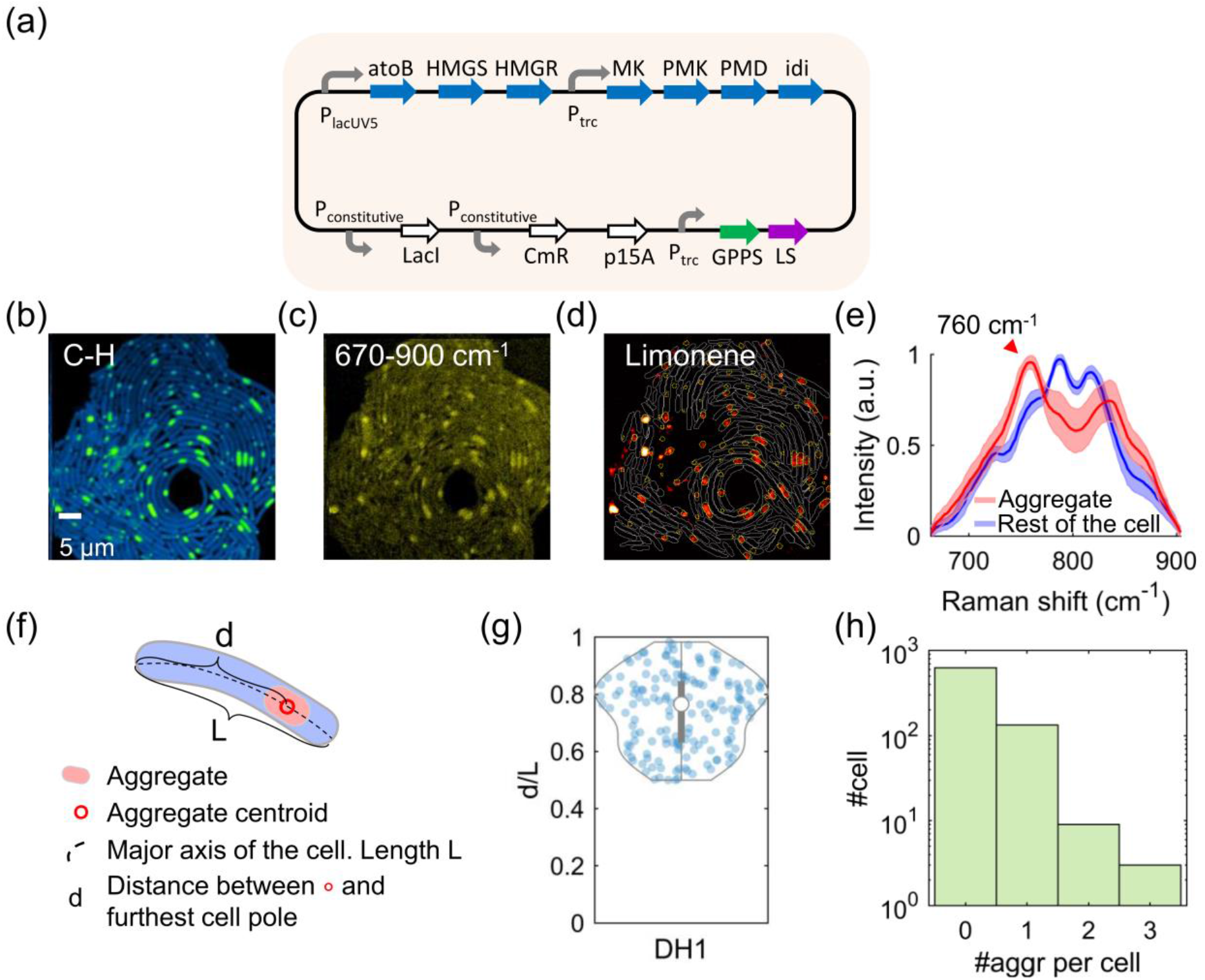
Characterization of limonene production in one-plasmid limonene-producing *E. coli*. (a) Genetic design for strains with one-plasmid system. (b-c) Spectrally summed hSRS images of DH1 one-plasmid strain. (d) Chemical maps of limonene channel of the same field of view in (b). (e) Averaged spectra of aggregates and rest of the cell (670 – 900 cm^−1^). Shaded error bar: std. (f) Diagram of a cell with an intracellular aggregate. (g) Violin plot with values of d/L for DH1 one-plasmid strain. (h) Histogram of aggregate counts per cell for DH1 one-plasmid strain.

HSRS imaging further allowed us to study the intracellular location of these aggregates as well as cell-to-cell variation. By extracting the central line along the major axis of each aggregate-containing cell, the position of the aggregate was determined as the pixel with the maximal intensity along the central line (**Figure 5f**). The scatter plot of the relative location of the aggregate (defined as d/L) is shown in **Figure 5g**, and 48.5% of aggregates resided between 70% to 90% of the cell length for DH1 strains. We also found that only 18.9% of the cells contain aggregates (**Figure 5h**). Similar results were observed in the BW25113 strain (**Figure S6g,h**). Cell-to-cell variation in limonene biosynthesis was also quantified using the average and total limonene content per aggregate (**Figure S7**). Interestingly, a larger aggregate tends to be accompanied by higher average limonene content (**Figure S7a**). Moreover, the scatter plot of aggregate size versus average limonene intensity was bounded by a constant upper threshold and an asymptotic lower threshold. In addition, by comparing the spectra of these aggregates and limonene solution in dodecane, the local concentration of limonene was estimated to be around tens to one hundred mM (**Figure S2**). Further evidence revealing the biosynthetic function of these proteins is shown in the next section.

### Intracellular limonene is co-localized with limonene synthase

To further validate our observation, simultaneous imaging of both limonene and LS, the final enzyme in the pathway that synthesizes it, was carried out. For this purpose, green fluorescence protein (GFP) was translationally fused to LS. Notably, limonene was observed in the strain with GFP-LS fusion (**Figure 6a,b**). Co-registered images of SRS and two-photon fluorescence of both BW25113 with two-plasmid system and DH1 with one-plasmid system are shown in **Figure 6c,d**, and **Figure 6f,g**, respectively. Limonene maps generated from SRS images are overlaid with the contour of the aggregates in the fluorescence channel, exhibiting the colocalization of limonene with GFP-fused LS (**Figure 6e,h**). Co-occurrence between limonene and limonene synthase was quantified by Mander’s colocalization coefficient (MCC) (39). MCC is defined as the summed limonene intensity of the colocalized pixels over the summed limonene intensity of all pixels (detailed definition in the Methods section). Higher MCC values indicate a higher colocalization level. BW25113 with two-plasmid system (**Figure 6e**) has an MCC of 0.79, and DH1 with one-plasmid system (**Figure 6h**) has an MCC of 0.71. This quantification confirms that these aggregates encompass both enzyme and limonene, with the potential function of limonene synthesis and storage. Together, our observation of colocalization of limonene and its synthesis enzyme verifies the formation of limonene synthesis metabolon inside the engineered *E. coli* strains.

**Figure 6.**
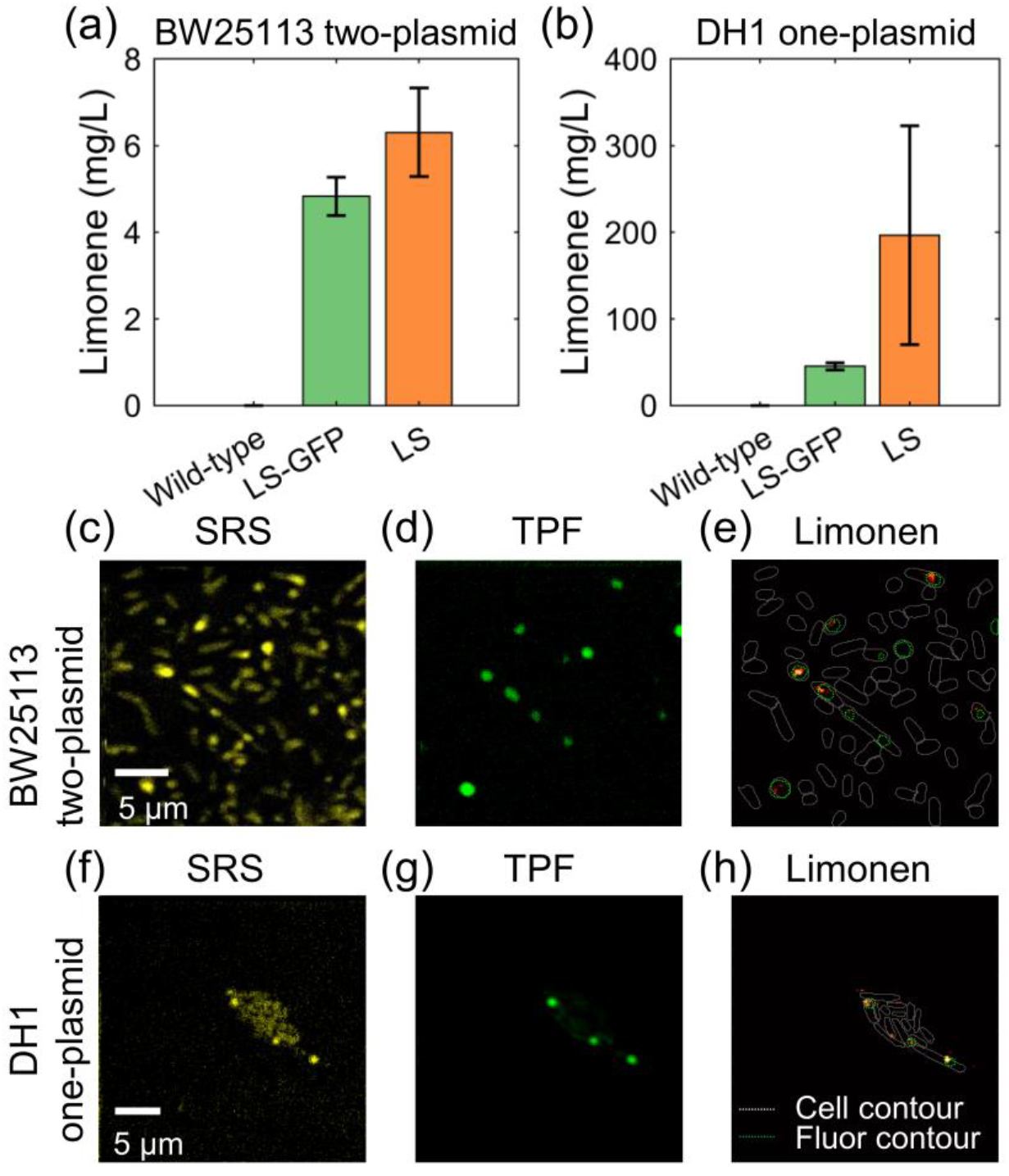
SRS and two-photon fluorescence (TPF) imaging unveil co-localization of limonene and limonene synthase. (a) GC-MS measurement of microbial production of limonene in BW25113 two-plasmid strain. Error bar: std. (b) GC-MS measurement of microbial production of limonene in DH1 one-plasmid strain. Error bar: std. (c) Spectrally summed hSRS images of BW25113 two-plasmid strain (670-900 cm^−1^ region). (d) Two-photon fluorescence image of the same FOV in (a). (e) Limonene map of the same FOV in (a-b). (f-h) The same order as (a-c). DH1 one-plasmid strain.

## Discussion

Limonene is a high-value chemical with various applications in the industry. A thorough understanding of the limonene biosynthesis process in the host microorganism is desired to guide metabolic engineering and potentially enhance the limonene yield. Our study here revealed the formation of limonene-rich aggregates in engineered *E. coli* strains with a limonene synthesis pathway. Quantification of the chemical content of these aggregates was achieved by spectral unmixing of hSRS images of spectral region 670 – 900 cm^−1^. Furthermore, taking advantage of simultaneous hSRS and TPF imaging, colocalization of limonene and limonene synthase was observed in the engineered *E. coli*. Two biological systems (one-plasmid DH1 strain and two-plasmid BW25113 strain) were tested and both exhibited aggregates in variants with a full limonene pathway. These data offer direct evidence of metabolite channeling and metabolon formation in these engineered strains.

Intracellular organization achieved through compartmentalization is key to cell metabolism. Consisting of sequential enzymes and metabolites, metabolon offers intracellular spatial organization in both eukaryotic (40, 41) and prokaryotic cells (42), bringing multiple advantages in terms of biosynthesis efficiency, cell growth, and survival (1, 43, 44). Both natural and engineered systems could take advantage of cellular localization (45, 46). Benefiting from the multimodal capabilities of our imaging system, this study suggests that these intracellular aggregates are limonene synthesis metabolons. First, limonene was shown enriched inside aggregates (**Figure 2–5**). In the DH1 one-plasmid strain, limonene concentration of aggregates is around tens to one hundred mM, which is one to two order magnitude higher than the limonene concentration measured by GC-MS (**Figure 5e**, **Figure S2a**). Moreover, larger aggregates were accompanied by higher limonene intensities, indicating the continuous limonene deposition in these regions (**Figure S7a**). Second, proteins were found existing inside the limonene-rich aggregates by comparing the C-H stretching spectra of aggregates with a standard protein sample BSA (**Figure 2g,h**). Considering the hydrophobicity of limonene and its precursors, these proteins may offer protection of these metabolites from the hydrophilic environment (5). Another advantage of these coexisting proteins is preventing the diffusion of reactive or potentially toxic intermediates, which potentially increases the reaction efficiency and reduces the metabolic cross-talk (47). Last, the enrichment of GFP-fused limonene synthase inside these aggregates suggests active limonene synthesis at these spots (**Figure 6**). This provides substantial evidence of the synthetic function of the proteins coexisting with limonene. Collectively, these data confirmed both the chemical composition and synthetic function of these intracellular aggregates, documenting a new case of metabolon formation in limonene-producing *E. coli*.

HSRS imaging also offers an interesting insight into the cell-to-cell variation of these limonene-producing cells. For all the strains with the full limonene synthesis pathway in this study, heterogeneity in terms of intracellular localization and limonene content were observed. Most of the cells contain zero or one aggregate, suggesting the sparsity of the high-producers in the whole population (**Figures 5h, Figure S6h**). The histogram of average limonene content per aggregate showed about ten times variation (**Figure S7b**) within an isogenic population for both BW25113 and DH1 one-plasmid strains. We are also able to observe the spatial bias where aggregates were formed. Asymmetric cell partitioning might be one possible reason for the observed single-cell heterogeneity and the pole-biased accumulation in these limonene-producing *E. coli* (**Figure 5f,g, Figure S6g**) (48, 49).

We would also note a few limitations of our method in characterizing the proposed limonene synthesis metabolon. First, we assumed that at each single pixel, the chemical composition is sparse. By using three standard reference spectra: pure limonene, the wild-type strain, and the background (of each hSRS image stack), the potential Raman signal contribution from the precursors was forced to go to the cell body channel. To discriminate limonene and three precursors (IPP, DMAPP, and GPP), a broadband coherent Raman imaging system could be applied (50, 51). By collecting a wider Raman spectral window, more metabolites could be quantified simultaneously, providing direct evidence of metabolite channeling (52). Second, limited by the limonene detection sensitivity (~21 mM) of our hSRS imaging system, limonene distribution in the non-aggregating area of the cell was not ruled out. Although most limonene is expected to exist in an aggregate form because of its low solubility in a hydrophilic environment, limonene efflux (53) is still of special interest and could be achieved by surface-enhanced Raman scattering (54). Third, time-lapse imaging of the limonene-producing *E. coli* was not achieved in this study due to laser toxicity (55). Machine learning-based methods offer a potential way to mitigate laser toxicity by overcoming tradeoffs between imaging speed, sensitivity, and specificity (56, 57).

In summary, our study provides direct visualization of the chemical composition and synthetic function of the limonene-rich aggregates inside engineered *E. coli*. This evidence of channeled limonene and its colocalization with limonene synthase is an important addition to our knowledge of the spatial control of limonene biosynthesis in engineered *E. coli*, and can offer conceivable guidance on metabolic engineering designs. Further investigations of localized limonene biosynthesis may include broadband coherent Raman imaging and time-lapse SRS imaging of live *E. coli* cells. With the live-cell imaging capacity, integration of our imaging platform with downstream cell sorting (58, 59) is expected.

## Materials and Methods

### Sample preparation

The *E. coli* strains in this study were derived from the BW25113 and DH1 strains. These wild-type strains were transformed with plasmids expressing the heterologous pathways for limonene production. For the two-plasmid system, the limonene synthesis pathway is split among two plasmids. First, the mevalonate pathway was expressed using plasmid pJBEI-3085 from Taek Soon Lee (Addgene #87950). Second, GPPS was expressed from a plasmid derived from pJBEI-3933 (60), but where we replaced pinene synthase with limonene synthase using golden gate cloning. pJBEI-3933 was a gift from Jay D. Keasling. For the one-plasmid system, the transformed plasmid was pJBEI-6409 from Taek Soon Lee (Addgene #47048) or variants of this plasmid.

Cell cultures were inoculated in Luria Bertani (LB) medium with appropriate antibiotics for plasmid maintenance (two-plasmid system: 25 μg/mL chloramphenicol and 100 μg/mL carbenicillin; one-plasmid system: 25 “μg/mL” chloramphenicol). On the following day, the cell culture was refreshed in 3 mL of M9 media (M9 salts, 2mM MgSO_4_, 100 μl CaCl_2_, 0.2% casaminos acids, 340 mg/L thiamine) supplemented with 20 g/L glucose and appropriate antibiotics. Limonene was induced by isopropyl β-d-1-thiogalactopyranoside (IPTG) when OD600 (optical density at 600 nm) reached ~0.8. The induction level was 100 μM for two-plasmid strains and 50 μM IPTG for one-plasmid strains if not otherwise specified. SRS images were taken after about 72 hours for two-plasmid strains and 24 hours for one-plasmid strains (except for the one-plasmid BW25113 strain in Figure 2, which was imaged after 24 hours). For SRS imaging of cells grown in M9 culture medium, 2-4 μL of cell culture was placed on a poly-l-lysine coated coverslip and sandwiched with another coverslip on the top. For SRS imaging of cells grown on M9 media agarose pads, 3 μL of cell culture was placed on a 3% agarose pad and sandwiched with another coverslip.

### Gas-chromatography mass-spectrometry (GC-MS)

Limonene samples were analyzed with an Agilent GC-MS 6890N equipped with an MS detector for up to 800 m/z. Helium was used as a carrier gas at a constant flow rate of 1 ml/min in an Agilent 222-5532LTM column. The inlet temperature was set to 300°C. The oven temperature was held at 50°C for 30 seconds, ramped up to 150°C at a rate of 25°C/min, and then further ramped to 250°C at a rate of 40°C/min. The results were analyzed using the MSD Productivity ChemStation (E.02.02.1431). An internal standard of α-pinene was used as a reference to calculate limonene concentrations.

### Ultrafast-tuning spectroscopic SRS microscope

SRS images were acquired using a lab-built SRS microscope (**Figure S1**) previously reported in ref (22). A polygon scanner (Lincoln SA24, Cambridge Technology) scanned the Stokes beam onto a blazed grating (GR50-0310, Thorlabs). As a reflective wedge, the grating introduces a continuous temporal delay between the pump and the retroreflected Stokes beam. Both the pump and Stokes beams were chirped with high-dispersion glass (SF57, 90 cm in length for the Stokes beam and 75 cm in length for the pump beam). For all experiments, the power on the sample was 20 mW for 966 nm, 14 mW for 800 nm, and 75 mW for 1040 nm, if not otherwise noted. The microscope was equipped with a 60x water immersion objective (NA = 1.2, UPlan-Apo/IR, Olympus). The SRS signal was then captured by a photodiode with a custom-built resonant circuit and extracted by a lock-in amplifier (UHFLI, Zurich Instrument). The same setup was used for two-photon fluorescence imaging with 20 mW for 966 nm. After filtering the excitation beam following the interaction with the sample, a photomultiplier tube (H7422-40, Hamamatsu) was used to measure the two-photon fluorescence signal.

### Linear unmixing of hyperspectral SRS signals

For each hyperspectral SRS image stack captured, a pixel-wise spectral unmixing was performed by the built-in MATLAB function LASSO (Least absolute shrinkage and selection operator). For every single pixel, LASSO decomposes its spectrum into the combination of several pure chemicals: *D* = *CS* + *E*. *D* is the detected signal 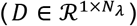. *C* is the decomposed concentration 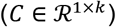. *S* is the measured spectral profiles of pure chemicals 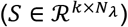. *E* is the residual term. *N_λ_* and *k* are the number of frames and the number of pure components, respectively. To get the optimal solution for this least-square fitting problem, L_1_-norm regularization was applied and the optimization problem is formulated as: 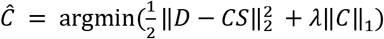. *λ* is a positive real number that denotes the regularizer penalty level. The same *λ* was chosen for a set of data recorded in the same imaging and digitizing conditions. Three spectral profiles were chosen as *S* including pure limonene, wild-type strain, and background (of each SRS image stack). The averaged Raman spectrum from wild-type was used as a cell background signal as it does neither show limonene content in GC-MS measurement nor Raman peak unique to limonene in SRS spectra. Averaged spectrum of the non-cell area of each SRS image stack was used as a reference to eliminate the background contribution from measurement.

### Image segmentation and analysis

For the hyperspectral SRS images, cell and aggregate segmentation was performed using the pixel classification + object classification workflow in ilastik (61). For cells grown into the clustered colony, an additional step of manual correction from Schnitzcells was applied in MATLAB to ensure high segmentation accuracy (62). Using these single-cell segmentation maps, quantification of single-cell features was performed by home-built MATLAB (MathWorks) scripts. Five categories of cell features were quantified: cell feature (area, major axis length), aggregate feature (area, major axis length), intracellular location of aggregate (distance from the aggregate centroid to the furthest cell pole), SRS image feature (SRS intensity at limonene peak 760 cm^−1^, spectrally summed SRS intensity at C-H and fingerprint window), chemical profile (limonene channel, cell body channel). For quantification of colocalization of limonene and limonene synthase, Mander’s colocalization coefficient was defined as 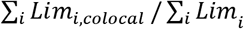, where *Lim_i,colocal_* = *Lim_i_* if *LimSyn* ≥ *threshold*, and *Lim_i,colocal_* = 0 if *LimSyn* < *threshold*.

## Acknowledgments

This work is supported by DOE Grant BER DE-SC0019387 to MJD, JXC, WW, and R35GM136223 and R01EB032391 to JXC. We thank Jean-Baptiste Lugagne and Eric South for helpful discussion on this project. We thank Boston University Chemical Instrumentation Center for their assistance with GC-MS experiments.

