## Supplementary figures for "Visualization of a Limonene Synthesis Metabolon inside Living Bacteria by Hyperspectral SRS Microscopy"

**This PDF file includes:**

Figures S1 to S7

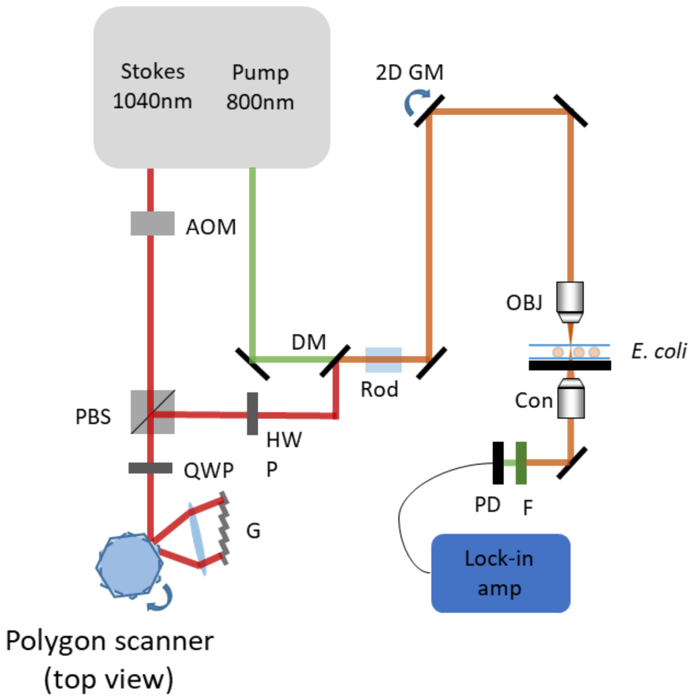


**Supplementary Figure 1. Schematic diagram of ultrafast spectroscopic SRS**

AOM, acousto-optic modulator; C, condenser; F, filter; GM, galvo mirror; HWP, half-wave plate; L, lens; LIA, lock-in amplifier; OBJ, objective; PBS, polarizing beam splitter; PD, photodiode; PS, polygon scanner; QWP, quarter-wave plate.

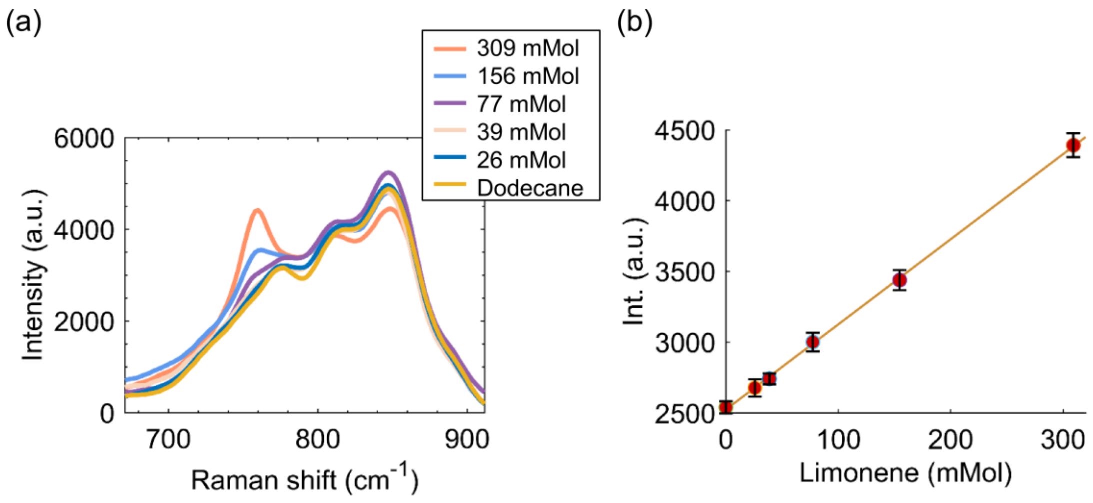


**Supplementary Figure 2. SRS spectra of limonene solution in dodecane with different concentrations**

(a) SRS spectra of limonene at different concentrations. Acquisition speed: 0.89 seconds per frame, 128 frames per image stack.

(b) Linear dependence of SRS intensity (at 760 $cm^{-1}$) on limonene concentration. The error bars show the noise level on the images (Standard deviation of 10 batches. Each batch is composed of 3x3 pixels).

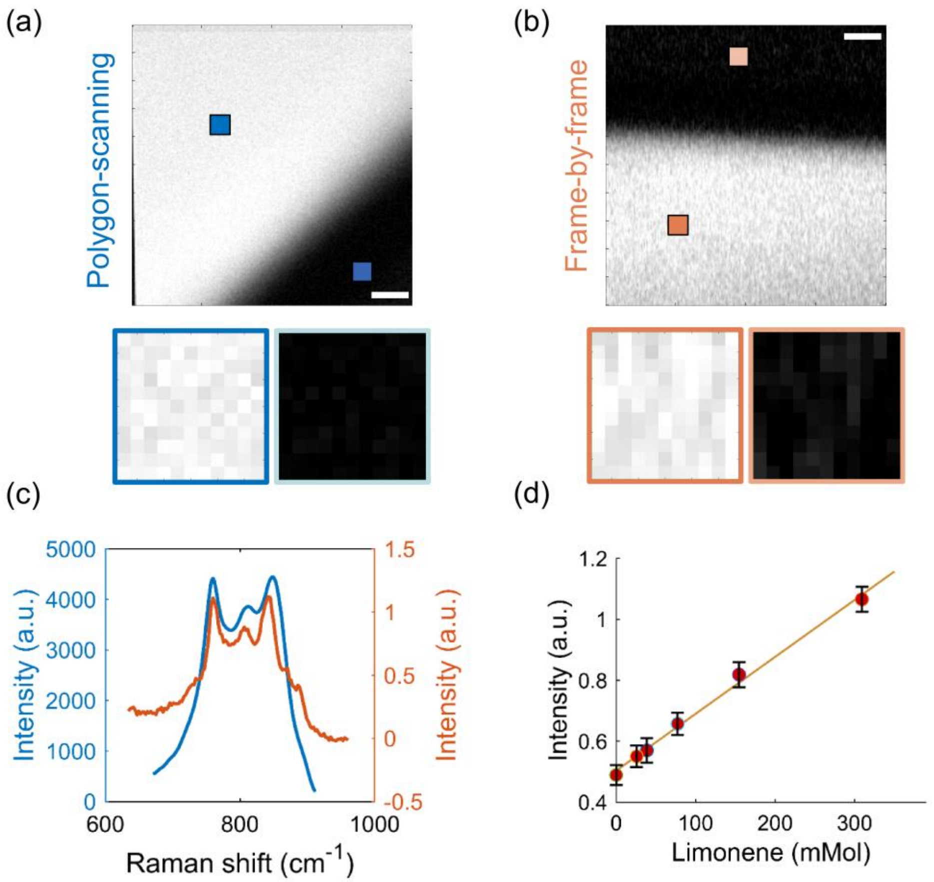


**Supplementary Figure 3. Comparison between polygon-scanning and frame-by-frame SRS**

(a) Single frame SRS at 760 cm-1 of 20x diluted limonene acquired in polygon-scanning SRS. Image contrast: [0 4414]. Scale bar: 10 µm.
(b) Single frame SRS at 760 cm-1 of 20x diluted limonene acquired in frame-by-frame SRS. This image was acquired using the same pump and Stokes power as polygon-scanning scheme. Image contras: [0 1.135]. Scale bar: 20 µm.
(c) Spectra of 20x diluted limonene from the squared area in (a) and (b). Blue: spectrum collected by polygon-scanning system; orange: spectrum collected by frame-by-frame system. The different signal scale in these two imaging schemes were due to different data acquisition strategies.
(d) Linear dependence of SRS intensity (at 760 cm-1) on limonene concentration in frame-by-frame scheme. These image stacks (200x200x128) were acquired using the same pump and Stokes as polygon-scanning scheme. The detection limit is ~30.0 mM.

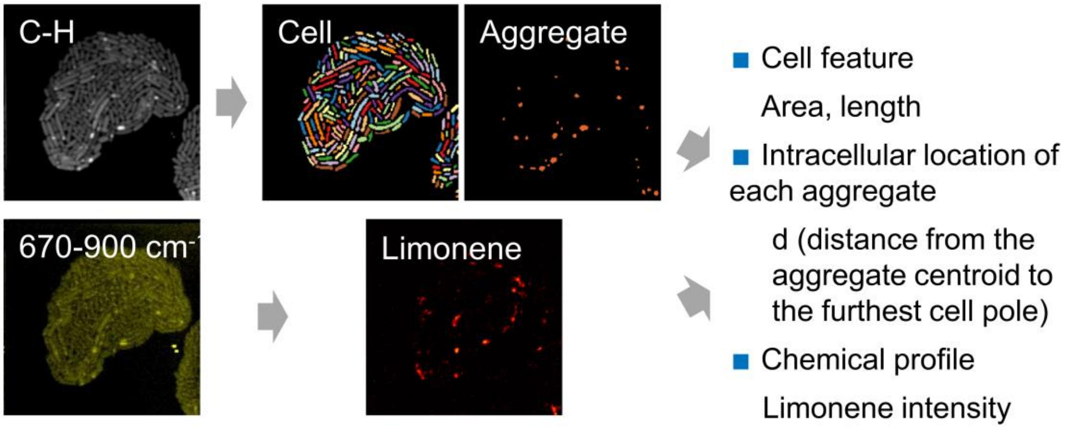


**Supplementary Figure 4. Single-cell and single-aggregate analysis pipeline**

Schematic representation of single-cell and single-aggregate analysis. The strain shown in this figure is *E. coli* DH1 one-plasmid strain.

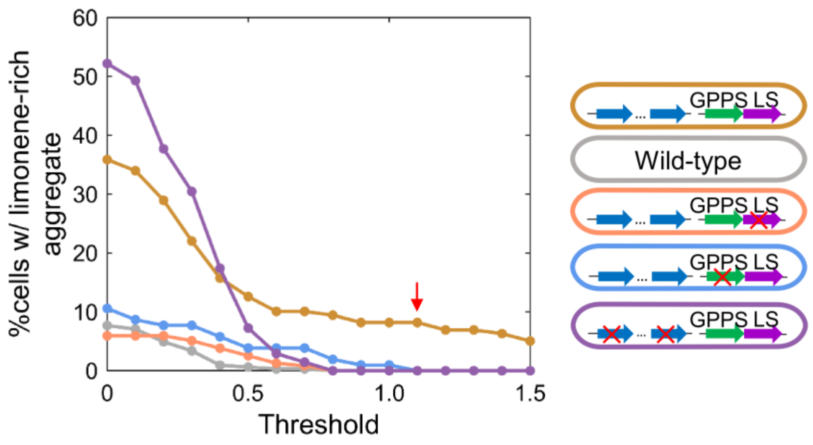


**Supplementary Figure 5. Determination of threshold of limonene-rich aggregate in *E. coli* with complete or incomplete limonene synthesis pathways.**

Different colors represent different genetic designs. The red arrow denotes the optimal threshold (1.1) of average limonene intensity per aggregate. At this threshold, all the other negative control strains do not have limonene-rich aggregates, and 16.9% of aggregates in the limonene-producing strain are limonene-rich.

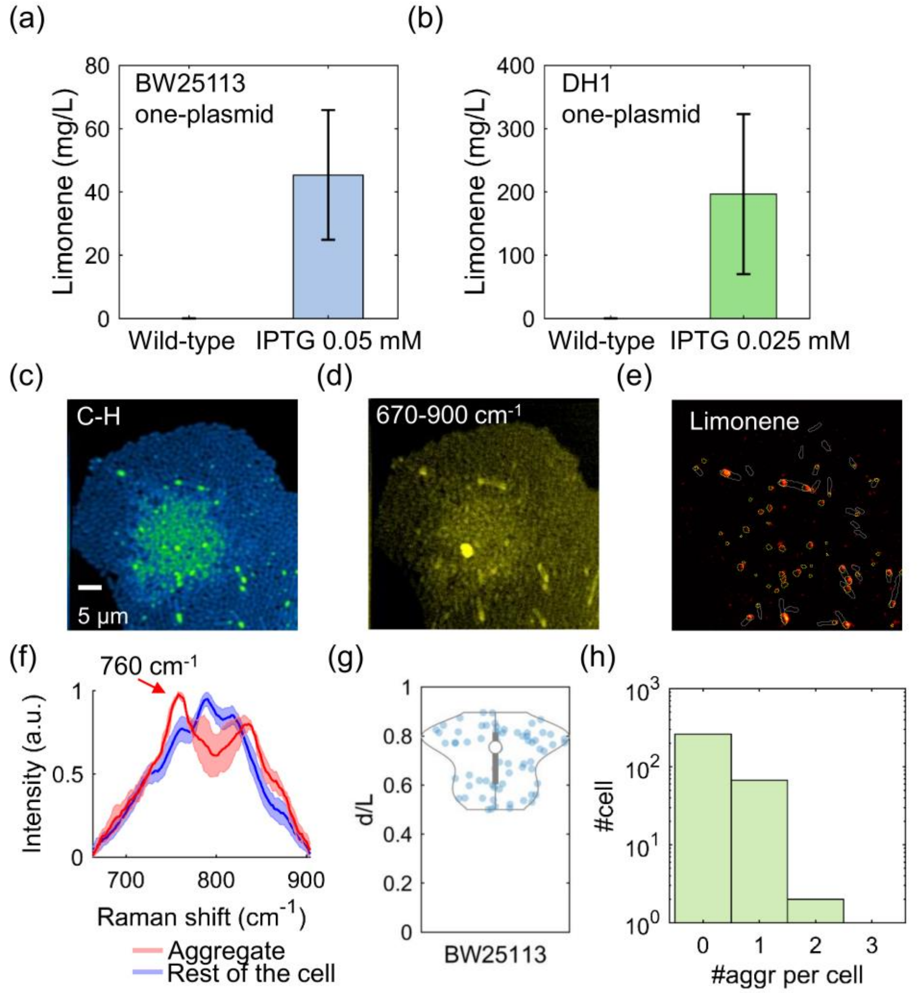


**Supplementary Figure 6. Characterization of limonene production in limonene-producing *E. coli***

(a) GC-MS measurement of microbial production of limonene in BW25113 one-plasmid strain. Error bar: std.
(b) GC-MS measurement of microbial production of limonene in DH1 one-plasmid strain. Error bar: std.
(c-d) Spectrally summed SRS images of BW25113 one-plasmid strain.
(e) Chemical map of limonene of the same field of view in (c-d).
(f) Averaged spectra (670 – 900 cm-1) of aggregate and rest of the cell. The spectrum of the aggregate shows the distinct Raman peak at 760 cm-1 unique to limonene. Shaded error bar: std.
(g) Violin plot with values of d/L for BW25113 one-plasmid strain. 56.3% of aggregates resided between 70% to 90% of the cell length.
(h) Histogram of aggregate counts per cell for BW25113 one-plasmid strain.

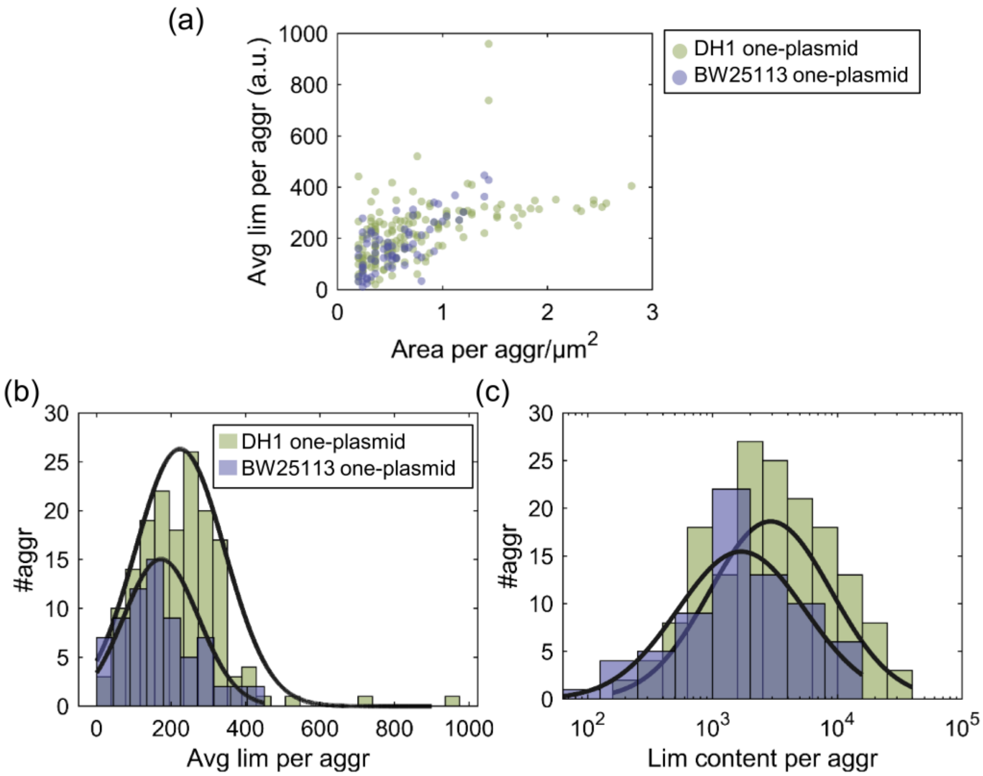


**Supplementary Figure 7. Characterization of aggregate formation in one-plasmid limonene-producing *E. coli***

(a) Scatter plot of area per aggregate versus average limonene content per aggregate.
(b) Histogram of average limonene content per aggregate. Both of the histogram are fitted to a Gaussian distribution and DH1 has significantly higher limonene concentration per aggregate (BW25113 one-plasmid: 173 ± 100, 5th percentile 24, 95th percentile 361; DH1 one-plasmid: 224 ± 12, 5th percentile 60, 95th percentile 393; arbitrary units; p = 0.0014, Wilcoxon rank-sum test).
(c) Histogram of total limonene content per aggregate. Both of the histograms are fitted to a log-normal distribution. (BW25113 one-plasmid: 5th percentile 156, 95th percentile 13620; DH1 one-plasmid: 5th percentile 60, 95th percentile 18178; arbitrary units)
